# Tumor relapse-free survival prognosis related consistency between cancer tissue and adjacent normal tissue in drug repurposing for solid tumor via connectivity map

**DOI:** 10.1101/2024.01.03.573997

**Authors:** Mingyue Hao, Dandan Li, Yuanyuan Qiao, Ming Xiong, Jun Li, Wei Ma

## Abstract

Traditional drug discovery encounters challenges, including high costs, time-intensive processes, and inherent risks. Drug repurposing emerges as a compelling alternative strategy to identify new indications for investigational or approved drugs, circumventing these obstacles. Among the various drug repurposing methods, the Disease-specific Signature-based Connectivity Map (Cmap) approach is widely utilized. However, the commonly employed method for constructing disease-specific signatures, known as Differentially Expressed Genes (DEG), faces issues related to inconsistencies between dysregulated genes and the prognosis of genes in tumor tissue, as well as discrepancies in prognosis genes between tumor and normal tissues.

In this study, we propose a novel approach, Prognosis Consistency Scoring (PCS), aimed at addressing these inconsistencies. PCS measures the consistency of gene prognosis between tumor and normal tissues by combining the Recurrence-Free Survival (RFS) prognosis power of genes in both contexts. Disease-specific signatures are then constructed based on PCS, and drug repurposing is performed using the Cmap and Lincs Unified Environment (CLUE). Validation of predicted drugs is conducted using data from DrugBank and ClinicalTrials databases.

Our findings reveal that the aforementioned inconsistencies are pervasive. Compared to signatures based on DEGs, PCS-based signatures exhibit superior performance, identifying more drugs with higher prediction accuracy, as confirmed by DrugBank annotations. Notably, a significant proportion of predicted drugs without corresponding indications were subsequently validated in the ClinicalTrials database. Additionally, PCS-based signatures demonstrate elevated disease specificity and association with Drug Related Gene (DRG).

## Introduction

Cancer poses a significant public health challenge, with an estimated 19.3 million new diagnoses and 10 million deaths reported in 2020[1]. Despite the discovery and utilization of numerous anti-cancer drugs, their efficacy is often compromised due to various factors, including toxicity, limited bioavailability, and drug resistance, attributable to the inherent heterogeneity and complexity of cancer[2]. Consequently, there is a pressing need to identify anti-cancer drugs exhibiting enhanced efficacy and reduced toxicity to enhance clinical outcomes. Conventional drug research and discovery encompass a series of stages, including target selection, compound optimization, safety analysis in animal models, and clinical studies (phases I–III). However, this traditional approach encounters significant bottlenecks marked by high costs, time-intensive procedures, and inherent risks[3, 4]. In response to these challenges, drug repurposing has emerged as a popular and effective alternative strategy in recent years. This innovative approach seeks to identify new indications for investigational or approved drugs, extending beyond their original medical scope, thereby offering a promising avenue to overcome the aforementioned obstacles[4–6]. Common methodologies for computational drug repurposing include machine learning-based approaches[7–9], network-based approaches[10–13], text mining-based approaches[14–16], semantics inference-based approaches[17, 18], sequence-based approaches[19], structure-based approaches[20–22], and signature-based drug repurposing approaches[3, 4]. Among these methods, signature-based approaches have rapidly evolved with the accumulation of big data in life sciences, such as the growing availability of gene expression data[3, 4]. The Connectivity Map (Cmap) is the prototypical drug repurposing approach in signature-based methodologies, and Cmap-based approaches dominate this category[3, 23].

Signature-based approaches necessitate extensive gene expression response profiles affected by chemical compounds, including approved drugs, and gene overexpression or knockdown. These datasets offer insights into the perturbation effects of chemical compounds or genes on the transcriptomics level of cell lines. The two primary data resources for this purpose are Cmap1 and Cmap2[4, 23–27]. The Cmap was introduced by Lamb *et al* in 2006[24]. The Cmap1 had two versions, build 1 and build 2[4]. The initial Cmap1 (build 1) database contained 564 gene expression profiles generated by treating 164 different approved small molecule drugs in five human reference cell lines at 42 concentrations and two time points[24]. The updated version, Cmap1 build 2, includes over 7,000 gene expression profiles associated with 1,309 approved drugs, 156 concentrations, and the same cell lines[3, 4]. The subsequent iteration, the next-generation version, Cmap2 or Library of Integrated Network-Based Cellular Signatures L1000 (LINCS-L1000), was developed by the same team in 2014[28]. It was completed in 2017 and generated 1,319,138 L100 expression profiles from 42,080 perturbagens, including 19,811 small molecule compounds, 18,493 shRNAs, 3,462 cDNAs, and 314 biologics[26].

The fundamental principle of Cmap-based drug repurposing involves comparing a disease-specific gene signature of transcriptome perturbation to Cmap reference profiles. Compounds that can reverse the disease-specific gene expression perturbation signature are prioritized as candidate drugs for treatment[24, 26]. Differential expressed genes (DEGs) serve as signatures in the Cmap reference database[24, 26], with DEGs or DEGs-based approaches being the most widely used methods for disease-specific signatures[4, 29–34]. Additionally, other methods exist, such as utilizing event-free survival via Cox regression to identify prognosis-related gene signatures[35], exploring disease-related genes through genome-wide association studies (GWAS)[36], conducting transcriptome-wide association studies (TWAS)[37], and using hub genes in protein-protein interaction (PPI) networks as queried signatures [38]. Functional annotations databases, text mining, and other methods are also applied to prioritize disease risk genes and generate signatures[39, 40].

However, in the context of solid tumors, two critical questions arise: (1) Does the dysregulated direction of DEGs in carcinoma consistently promote tumor progression? (2) Do genes in solid tumors and adjacent normal tissues play the same role in tumor progression? To address these questions, we introduced a new method to generate disease-specific signatures in this study. To answer question 1, tumor relapse-free survival (RFS) analysis via Cox proportional hazards models measured the tumor prognosis roles of genes in tumor tissues. To address question 2, the tumor prognosis roles of genes in tumor and normal tissues were combined into a prognosis consistency score (PCS). Our results indicate that when PCS-based disease-specific signatures were employed to query Cmap and Lincs Unified Environment (CLUE), a webserver tool of Cmap2, the drug prediction power of PCS-based signatures surpassed that of DEGs-based signatures.

## Materials and method

### Data source

The preprocessed RNA-sequencing (RNA-seq) data, normalized by RSEM, and corresponding clinical information of solid tumors from The Cancer Genome Atlas (TCGA) database were obtained from UCSC Cancer Browser (UCSC Xena, https://xenabrowser.net/datapages/) on August 28, 2020. The length of human species genes used in the analysis was retrieved from the BioMart tool of the Ensembl database (Ensembl, https://asia.ensembl.org/index.html) on March 20, 2021. Relevant literature was identified and acquired from the PubMed database (https://pubmed.ncbi.nlm.nih.gov/) on November 4, 2023. Drug annotation information was retrieved from the DrugBank database (https://www.drugbank.com/) version 5.1.10, released on January 4, 2023, accessed on September 8, 2023. Clinical trials data were downloaded from ClinicalTrials.gov database (https://www.clinicaltrials.gov/) on September 12, 2023. Disease-related genes (DRG) and the corresponding Disease Gene Score (DGS) were obtained from the CURATED subset of the DisGeNET database (https://www.disgenet.org/) via the disgenet2r R package on September 29, 2023. CLUE (https://clue.io/) was utilized to query disease-specific signatures for identifying candidate drugs. The connection score between perturbagens was obtained from the CLUE data source (https://clue.io/data/TS), and single drug or compound signatures were downloaded via the CLUE tool (https://clue.io/command). The 978 landmark genes, 11,350 inferred genes, and 9,196 best inferred genes were acquired from the supplementary materials of the original Cmap2 paper, titled "A Next Generation Connectivity Map: L1000 platform and the first 1,000,000 profiles"[26].

### Tumor selection

The focus of this study was on RFS events and tumor-normal paired samples. Transcriptome data and clinical data were matched, and tumor-normal paired samples were identified. Tumors with more than 40 tumor-normal paired samples with RFS information were selected for analysis, resulting in the selection of six types of tumors.

### DEG and Cox regression analysis

The edgeR R package was employed to identify DEGs via fold change (FC) and false discovery rates (*FDR*) between paired tumor and normal samples using RSEM counts. RSEM normalized counts data and gene length information were used to convert the data into transcripts per million (TPM) expression data, followed by log2 transformation (log2(TPM+1)). Univariable Cox regression analysis was performed using the survival R package, utilizing z-score normalized log2(TPM+1) for each gene to identify RFS-related genes in paired tumor and normal samples.

### Drug repurposing and prediction based on the tumor-specific signature via PCS and DEG

For drug repurposing and prediction via CLUE, a disease-specific signature needed to be constructed. The CLUE query tool (https://clue.io/query) was utilized for this purpose. Query parameters included the Gene Expression (L1000) dataset and version 1.0. A maximum of 150 up-regulated genes and 150 down-regulated genes were uploaded. The analysis focused on the 978 landmark genes and 9,196 best inferred genes. For DEG, genes were ordered by the log2-transformed fold change (logFC), and the top 150 and bottom 150 genes were extracted as the signature for up-regulated and down-regulated genes, respectively. The PCS score was calculated according to equation 1.

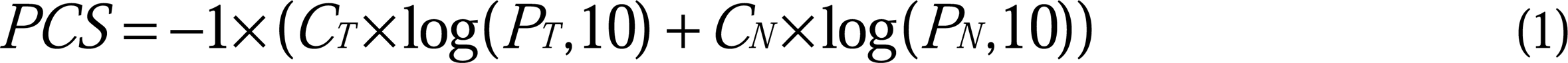

*C_T_* represents the Cox regression coefficient of tumor tissue, and *P_T_* signifies the *P*-value associated with the Cox regression coefficient of tumor tissue. Similarly, *C_N_*denotes the Cox regression coefficient of normal tissue, and *P_N_* represents the *P*-value linked to the Cox regression coefficient of normal tissue. Consequently, for the PCS, genes were ranked based on PCS, and the top 150 genes were submitted as the down-regulated gene list, while the bottom 150 genes were submitted as the up-regulated gene list. Upon completion of the query, the results were downloaded through the query history.

### Drug combination order analysis

For the identified candidate drug list, their signatures were acquired from the CLUE command tool (https://clue.io/command), and the summary signature was computed according to the method outlined at https://clue.io/connectopedia/replicate_collapse. Genes were then ranked by moderated z-score (MODZ, the weighted averages of z-score vectors calculated above), and the top 150 genes were extracted as the down-regulated gene signature, while the bottom 150 genes were extracted as the up-regulated gene signature. The intersection between the top 150 genes from PCS and the top 150 genes from the drug signature was calculated and defined as TopSet. Similarly, the intersection between the bottom 150 genes from PCS and the bottom 150 genes from the drug signature was calculated and defined as TailSet. Subsequently, the *m* drugs were ranked in descending order by the sum of the sizes of TopSet and TailSet, as per equations 2 and 3. The first drug was assigned rank 1. Next, genes from TopSet were removed in PCS top 150 genes, and genes from TailSet were removed in PCS tail 150 genes, following equations 4 and 5. This process was repeated for the remaining drugs until the last one. As a result, each drug was assigned a rank, and these ranks constituted their combination order.

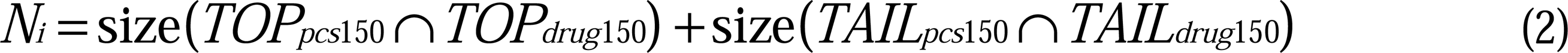

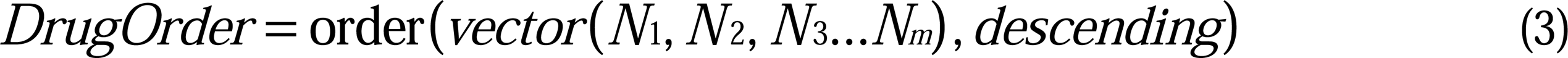

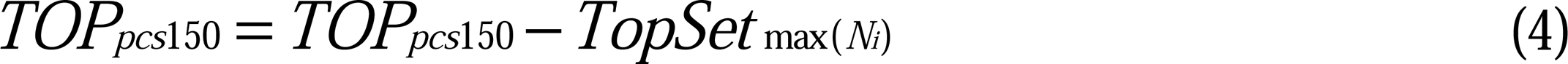

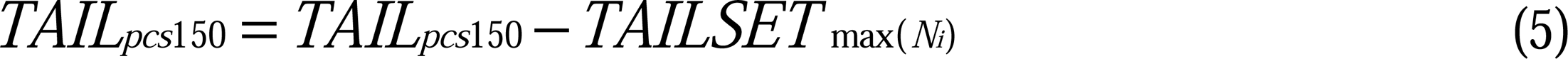

### Statistical analysis

Continuous variables underwent analysis using Student’s t-test, while categorical variables were assessed utilizing Fisher’s exact test. All statistical analyses and figure generation were conducted using R language version 4.1.1. R is an open-source environment for statistical computing and graphics, accessible at https://www.r-project.org/.

## Result

### Six types of solid tumor were selected for analysis

We gathered transcriptome data of tumors from the TCGA database. Solid tumors with more than 40 tumor-normal paired samples containing RFS information were chosen for analysis. Based on this criterion, six types of solid tumors, including breast invasive carcinoma (BRCA), head and neck squamous cell carcinoma (HNSC), liver hepatocellular carcinoma (LIHC), lung adenocarcinoma (LUAD), lung squamous cell carcinoma (LUSC), and thyroid carcinoma (THCA), were retained. The sample information for the six tumor types is summarized in Table 1.

**Table 1.** Sample information of the six types of tumor.

|  | BRCA | HNSC | LIHC | LUAD | LUSC | THCA |
| --- | --- | --- | --- | --- | --- | --- |
| **Tumor** | 1096 | 520 | 371 | 515 | 502 | 505 |
| **Normal** | 114 | 44 | 50 | 59 | 51 | 59 |
| **T-N P** | 114 | 43 | 50 | 58 | 51 | 59 |
| **T-N P RFS** | 114 | 43 | 41 | 53 | 40 | 58 |

T-N P: tumor-normal paired; T-N P RFS: tumor-normal paired sample with RFS information.

### Review of current signature-based drug repurposing approaches via Cmap

To survey current signature-based drug repurposing approaches via Cmap, we conducted a literature review of publications available before October 30, 2023, in the PubMed database, identifying a total of 313 relevant articles. After a comprehensive review, 190 articles were selected and summarized in Supplementary Table 1. The articles were annotated based on their signature-building approaches, the databases used, and their research objectives. We categorized the signature approaches into seven major groups and specific sub-approaches (see Table 2).

**Table 2.** Approaches used in the 190 literatures.

| Approach | Cmap1 | Cmap2 | Cmap1, 2 | Unknown | Sum |
| --- | --- | --- | --- | --- | --- |
| **DEG** | 61 | 74 | 11 | 0 | 146 |
| **Other based DEG** | 3 | 7 | 0 | 0 | 10 |
| **Disease related genes** | 4 | 2 | 2 | 1 | 9 |
| **DEG based** | 6 | 1 | 0 | 0 | 7 |
| **Target related genes** | 5 | 1 | 0 | 0 | 6 |
| **Prognosis related genes** | 1 | 3 | 1 | 0 | 5 |
| **DEG and others** | 3 | 0 | 0 | 0 | 3 |
| **Linear discriminant analysis** | 0 | 1 | 0 | 0 | 1 |
| **Network hub genes** | 0 | 1 | 0 | 0 | 1 |
| **Time-course trend genes** | 0 | 1 | 0 | 0 | 1 |
| **Transcriptomes correlations** | 1 | 0 | 0 | 0 | 1 |

The most frequently used approach was DEGs between disease samples and normal samples, accounting for 146 articles. Other DEG-based approaches, involving grouping based on methods other than tumor versus normal such as high-risk group versus low-risk group, constituted 10 articles[41–43]. Disease-related genes were used as signatures in 9 articles, sourced from text mining or databases[44–46]. Seven articles employed combinations of DEG-based approaches, where DEGs were first identified, and then additional methods were applied. For example, DEGs were called first, and then differential pathways were called based on DEGs, and the pathways were used to evaluate the similarity between diseases and drugs[47–49]. Target-related genes were utilized in 6 articles, where a disease target was confirmed, and the target-related genes were identified as signatures.[50–52]. Prognosis-related genes were used in 5 articles, employing the univariate Cox proportional hazards regression model[53–55]. Three articles combined DEG with other methods to build signatures, such as selecting genes satisfying both DEG and overall survival prognosis criteria[56, 57]. Additionally, four specific approaches were identified, including linear discriminant analysis, network hub genes, time-course trend genes, and transcriptome correlations. Linear discriminant analysis was employed to manage samples involving multiple conditions and time-points. In essence, it also computed DEGs, but these were referred to as meta-DEGs[58]. The methodology for time-course trend genes resembled that of linear discriminant analysis but was simpler in nature. It served to address samples spanning multiple time-points, concurrently calculating meta-DEGs[59]. In the case of transcriptome correlations, no specific signatures were computed. Instead, the entire transcriptome was treated as a signature, and measures of overall transcriptome similarity, such as Spearman correlation or cosine similarity, were calculated[60].

The 190 articles were further classified based on the databases used (Table 3). Eighty-four articles were Cmap1-based, 91 were Cmap2-based, and 14 utilized both. An annual analysis of the approaches and databases used in the articles is presented in Figure 1. The use of both Cmap1 and Cmap2 was classified under the Cmap2 group. Overall, the number of articles increased annually, with DEG being the most frequently employed method each year and in total, as depicted in the top-left of Figure 1.

**Table 3.** Databases used in the 190 literatures.

| Cmap | 11-16 | 17 | 18 | 19 | 20 | 21 | 22 | 23 | Sum |
| --- | --- | --- | --- | --- | --- | --- | --- | --- | --- |
| **1** | 26 | 8 | 3 | 11 | 4 | 13 | 12 | 7 | 84 |
| **2** | 0 | 1 | 3 | 6 | 16 | 19 | 26 | 20 | 91 |
| **1 and 2** | 0 | 0 | 2 | 5 | 2 | 1 | 3 | 1 | 14 |
| **UK** | 0 | 0 | 0 | 0 | 0 | 0 | 0 | 1 | 1 |

UK: unknown.

**Figure 1.**
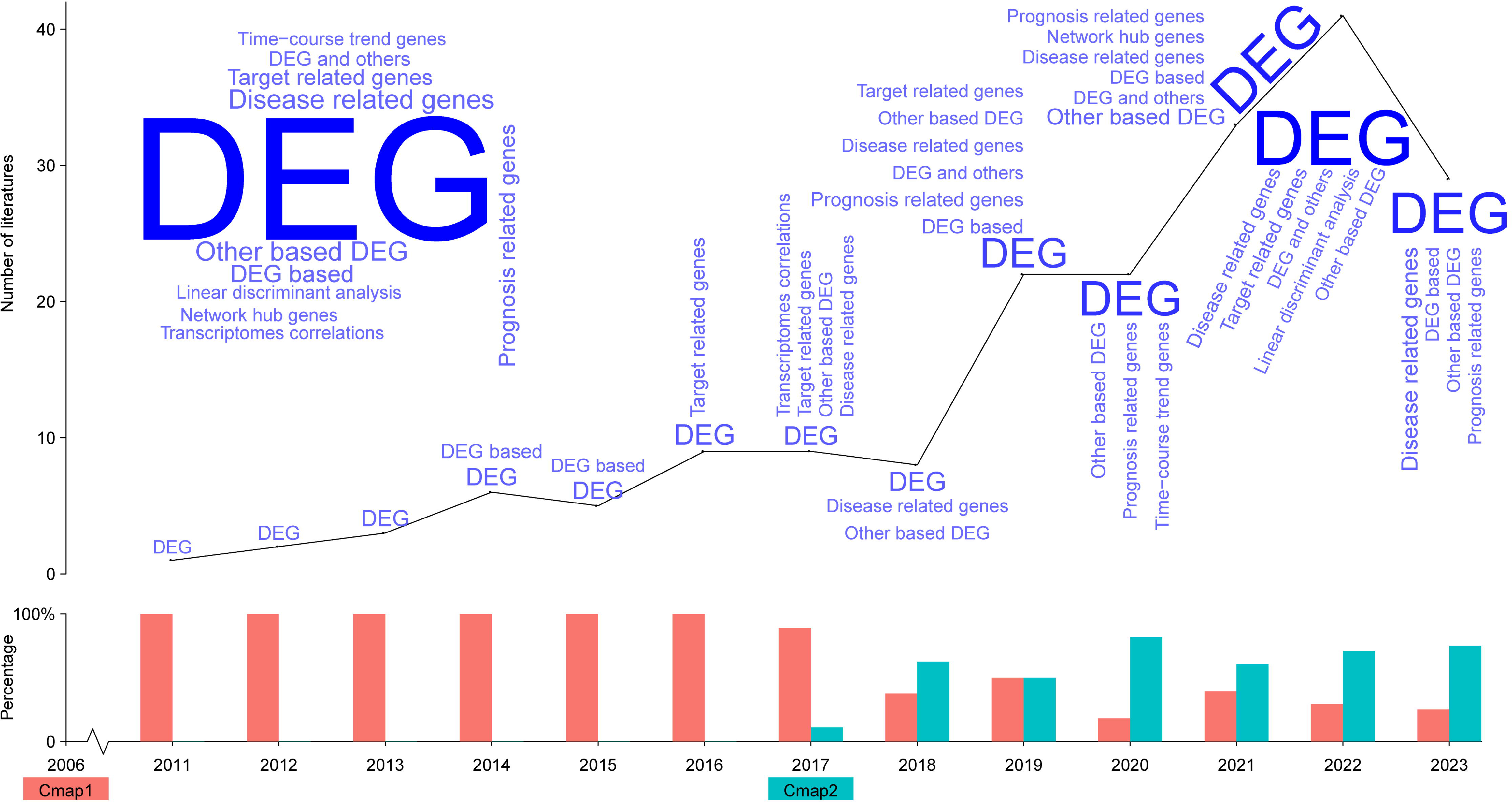
Signature building methods and used databases from 2011 to 2023. Top-left: Word cloud of all signature building methods.

### Inconsistency between dysregulation and RFS prognosis of genes

In the investigation across six types of solid tumors, we examined the concordance between differential expression and RFS prognosis of genes. Differential expression gene sets were calculated between tumor and normal tissues across tumor-normal paired samples, along with the identification of RFS prognosis-related genes in tumor samples paired with normal samples. Twelve cutoff values were set (DEG-*FDR*: 1, 0.9, 0.8, 0.7, 0.6, 0.5, 0.4, 0.3, 0.2, 0.1, 0.05, 0.01; DEG-abs(logFC): 0, 0.1, 0.2, 0.3, 0.4, 0.5, 0.6, 0.7, 0.8, 0.9, 1, 2; Cox-*P*-value: 1, 0.9, 0.8, 0.7, 0.6, 0.5, 0.4, 0.3, 0.2, 0.1, 0.05, 0.01) to explore the consistency between dysregulation and RFS prognosis. For each tumor type, four sets were generated: tumor up-regulated genes set (TU), tumor down-regulated genes set (TD), tumor RFS Cox positive coefficient set (TRFSP), and tumor RFS Cox negative coefficient set (TRFSN). Consistency genes were defined as the intersection set between TU and TRFSP and the intersection set between TD and TRFSN. Unconsistency genes were defined as the intersection set between TU and TRFSN and the intersection set between TD and TRFSP. The unconsistency ratio of dysregulation and RFS prognosis (UCRDP) was calculated as the ratio between the number of unconsistency genes and the number of genes in the union of the four intersection sets.

As depicted in Figure 2, in LUSC and THCA, UCRDP increased with stricter thresholds. A Venn diagram provided further details at a common cutoff (*FDR* = 0.05, *P*-value = 0.05, Log(FC) = ±1). The UCRDP in LUSC and THCA consistently exceeded 50%. Conversely, UCRDP decreased with stricter thresholds in BRCA, LIHC, LUAD, and HNSC. UCRDP maintained a range of 10% to 50% in BRCA, LIHC, LUAD, and THCA, except for the common cutoff point in LUAD. The last point in LIHC, LUAD, BRCA, and THCA was zero cause of limited genes. This analysis revealed that inconsistency between dysregulation and RFS prognosis was a prevalent phenomenon, exceeding consistency in certain tumor types. Thus, the direction of differential expression genes did not singularly indicate their role in tumor relapse progression.

**Figure 2.**
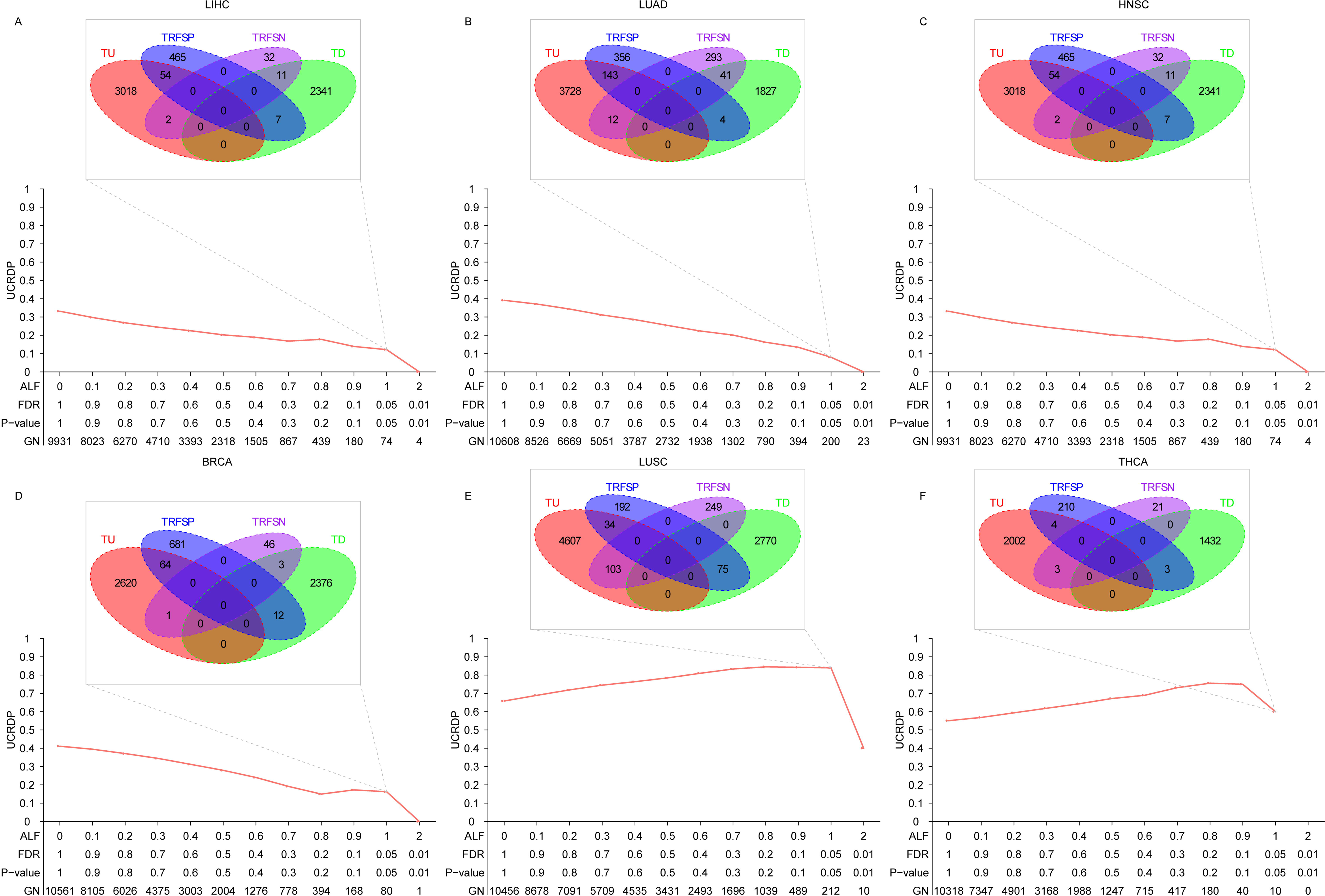
UCRDP in the six types of tumor. UCRDP: Unconsistency ratio of dysregulation and RFS prognosis; ALF: Absolute value of logFC; GN: Number of genes in the union of unconsistency genes and consistency genes.

### Inconsistency between RFS prognosis of genes in tumor and adjacent normal tissue

The study delved into the alignment between RFS prognosis of genes in the tumor and adjacent normal tissue across the six solid tumors using 12 cutoffs (Cox-*P*-value: 1, 0.9, 0.8, 0.7, 0.6, 0.5, 0.4, 0.3, 0.2, 0.1, 0.05, 0.01). For each tumor type, four sets were established, comprising tumor RFS Cox positive coefficient (TP), tumor Cox negative coefficient (TN), normal Cox positive coefficient (NP), and normal Cox negative coefficient (NN). Unconsistency genes were defined as the intersection between TP and NN, and the intersection between TN and NP. Consistency genes were defined as the intersection between TP and NP, and the intersection between TN and NN. The ratio of the number of unconsistency genes to the number of the union of unconsistency and consistency genes was termed unconsistency ratio between tumor RFS prognosis and normal RFS prognosis (UCRTPNP), serving as a metric to assess consistency.

As depicted in Figure 3, UCRTPNP exhibited a decreasing trend as the Cox-*P*-value reduced across all six types of solid tumors. The majority of UCRTPNP values remained below 50%, with a range of approximately 5% to 50%. While UCRTPNP generally demonstrated lower values compared to UCRDP, such observations were common. This analysis underscores the prevalent inconsistency between tumor and normal tissue in terms of RFS prognosis, highlighting the nuanced nature of gene behavior in these contexts.

**Figure 3.**
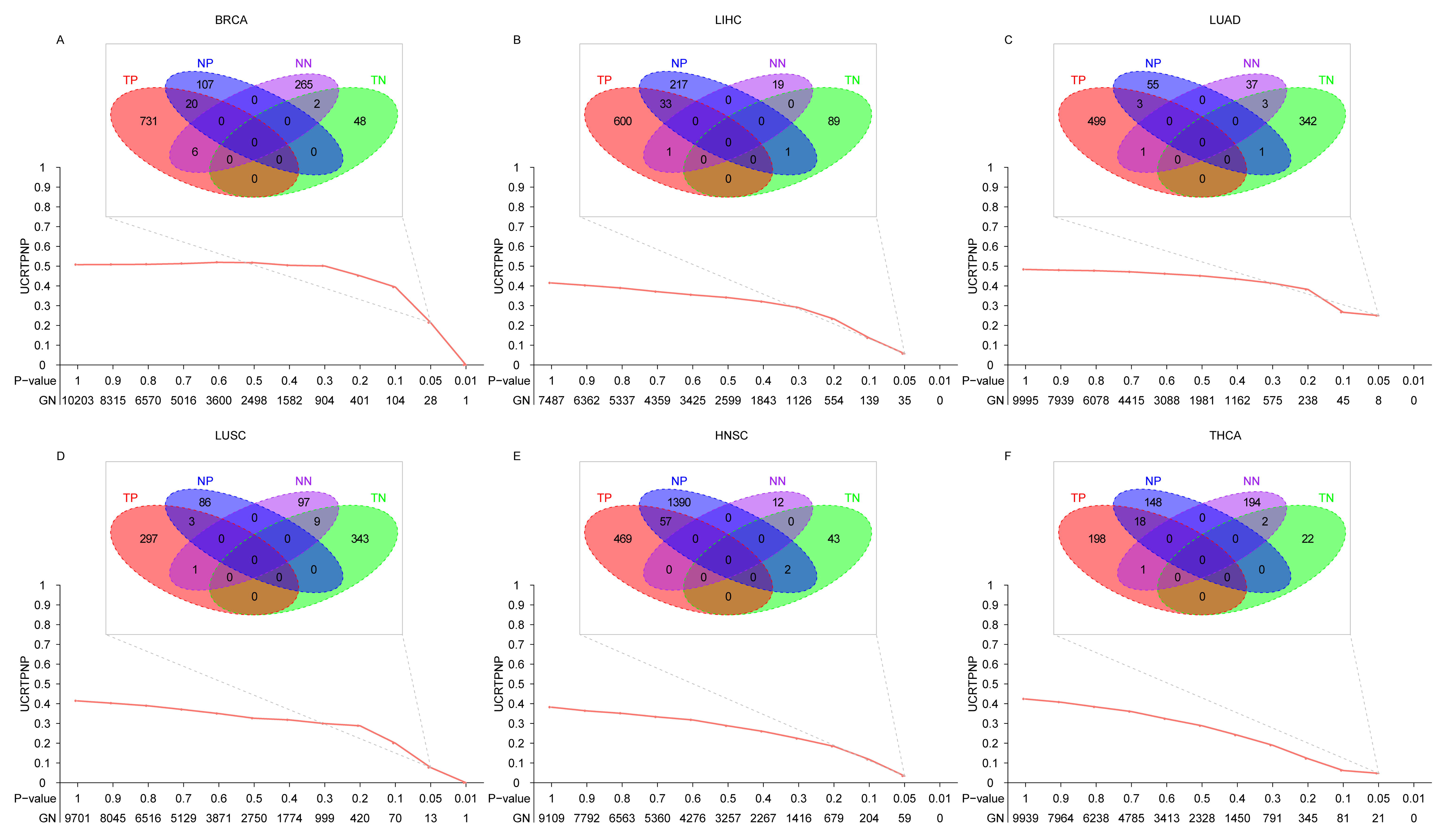
UCRTPNP in the six types of tumor. UCRTPNP: Unconsistency ratio between tumor RFS prognosis and normal RFS prognosis; GN: Number of the union of unconsistency and consistency genes.

### Drug repurposing and prediction power based on PCS signatures via CLUE

Cmap2, as the advanced iteration, assessed the transcriptome of 978 landmark genes, with an additional 11,350 genes inferred from this set. Within the inferred genes, 9,196 were identified as best inferred due to their high accuracy, while 2,154 genes had poor inference outcomes[26]. The disease-specific signature generation process involved utilizing the 978 landmark genes and the 9,196 best-inferred genes. For drug repurposing and prediction, the CLUE platform (https://clue.io/) was employed. Given CLUE’s requirement of no more than 150 genes in the up-genes or down-genes list, genes were ordered by LogFC or PCS, and the top 150 and tail 150 genes were extracted as the disease-specific signature. The top 150 genes were designated as the up-genes list, while the tail genes were assigned to the down-gene list. In the query results, compounds of touchstone were selected for further analysis. Candidate drugs were identified based on a high level of negative scores. The compounds were cross-referenced with the DrugBank database for verification of their status as drugs, including their indications and stage. Of the 2,427 compounds, 1,180 were validated in the DrugBank database.

Among the 1,180 drugs, 41 were found to have indications for breast cancer, with 35 being approved and 6 under other stages (Figure 4A and 4B). For PCS-based signature, 25 out of the 41 drugs received negative scores, 2 had zero scores, and 14 had positive scores (Figure 4A). Notably, 2 drugs scored lower than -90, 4 scored lower than -50, and 4 scored higher than 50 (Figure 4A). In contrast, DEGs-based signature only yielded 5 drugs with negative scores, 13 with zero scores, and 23 with positive scores (Figure 4B), and no drug scored lower than -90 or -50, while 2 scored higher than 90, and 6 scored more than 50 (Figure 4B). In BRCA, PCS-based signature outperformed DEGs-based signature.

**Figure 4.**
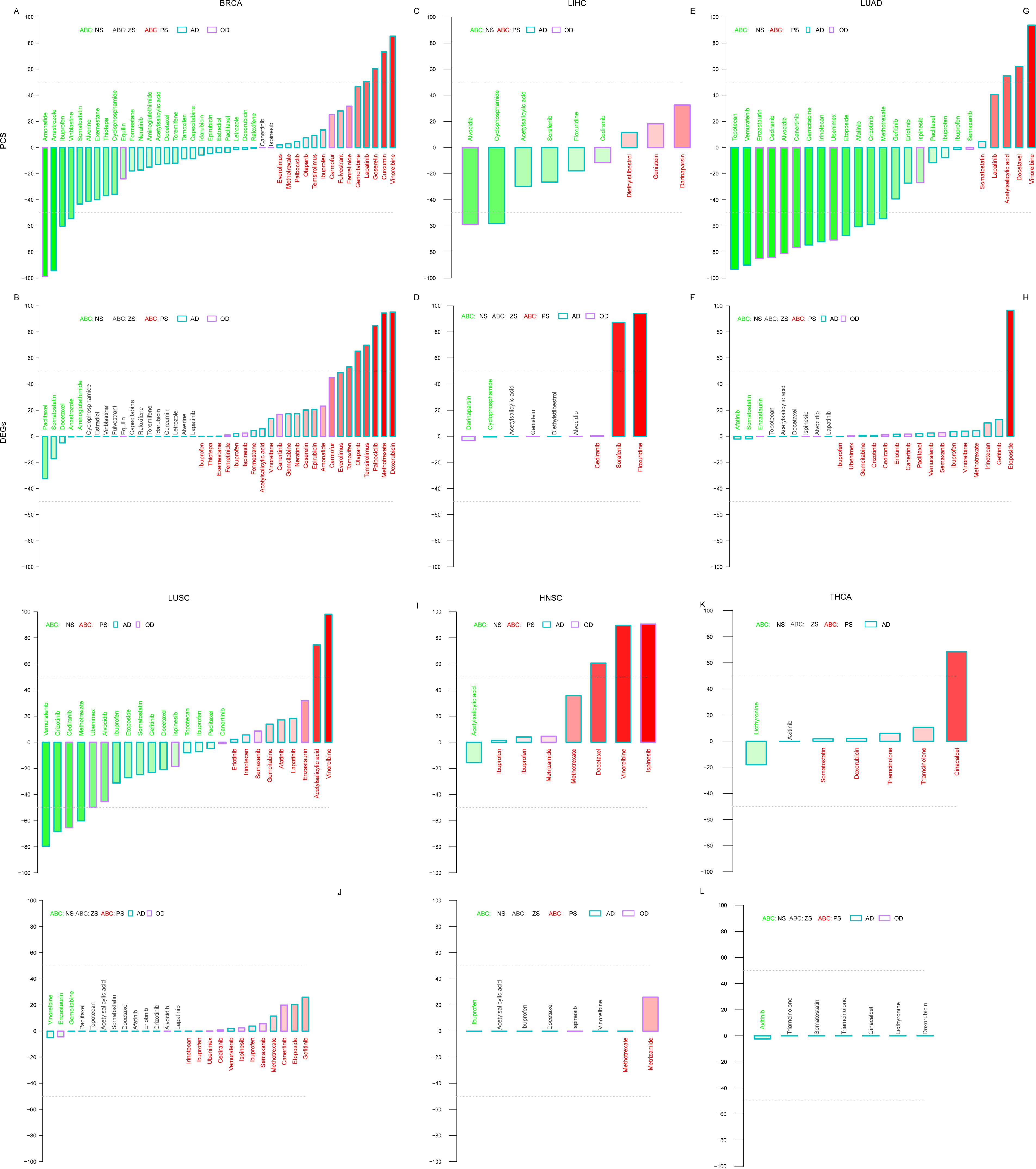
Drug repurposing prediction power of PCS and DEGs validated by DrugBank.

In LIHC, 9 drugs, including 5 approved and 4 in other stages, were found to have indications for liver cancer. For PCS-based signature, 6 drugs had negative scores, and 3 had positive scores (Figure 4C). Among the 6 negative score drugs, 2 scored lower than -50. In contrast, DEGs-based signature revealed 2 drugs with high positive scores, one even higher than 90, while other drugs had nearly zero scores (Figure 4D). In LIHC, PCS-based signature also outperformed DEGs-based signature. PCS identified more drugs (6) with indications, but without extremely low negative scores, whereas DEGs identified no drugs even two with high positive scores.

There were 25 drugs with indication of lung cancer. In LUAD, PCS-based signature indicated 20 drugs with negative scores and 5 with positive scores (Figure 4E). Among the 20 negative score drugs, 2 scored lower than -90, and 13 scored lower than -50. Among the 5 positive score drugs, 1 scored more than 90, and 3 scored more than 50. In contrast, DEGs-based signature identified nothing, even one with score higher than 90, and none of the other drugs scored higher than 20 or lower than -20 (Figure 4F). In LUAD, PCS demonstrated absolute predominance compared to DEGs.

In LUSC, PCS identified 16 drugs with negative scores and 9 with positive scores (Figure 4G). Among the 16 negative score drugs, 4 scored lower than -50, and among the 9 positive score drugs, 2 scored more than 50, with 1 scoring more than 90. In contrast, DEGs identified no drugs with scores lower than -50 (Figure 4H). Thus, in LUSC, PCS also outperformed DEGs.

In HNSC, 8 drugs, including 6 approved drugs and 2 in other stages, were identified to treat HNSC. For DEGs, no drugs scored more than 50 or lower than -50, while in PCS, 3 drugs scored more than 50, with 2 scoring more than 90 (Figure 4I and 4J). However, since the low negative drugs were targets treated as candidate drugs, PCS and DEGs performed similarly when only considering our interested targets, as both selected nothing. Nevertheless, PCS identified some high positive score drugs, indicating its opposite predicted results.

For THCA, 7 approved drugs were found. Similar to HNSC, PCS and DEGs selected none of the interested targets, but PCS predicted 1 opposite result (Figure 4K and 4L). Overall, considering all six types of tumors, PCS demonstrated better performance.

### Validation of predicted drugs via ClinicalTrials database

Following the query results, candidate drugs with low negative scores were identified for each of the six tumors. To substantiate these predictions, compounds with an absolute score above 50 were selected, and a search was conducted in the ClinicalTrials database for evidence of their efficacy against the six tumor types. Results were meticulously reviewed term by term on https://clinicaltrials.gov/, and findings were collected and summarized in Supplementary Table 2. When a drug undergoes multiple clinical trials for a specific type of tumor, affirmation of treatment efficacy in at least one trial implies a positive treatment effect for the tumor. Conversely, if all clinical trials unanimously indicate a lack of treatment effect, the drug is deemed ineffective for treating the tumor.

In BRCA, two drugs were identified for PCS (imatinib, -51.34; metformin, 55.28), and one for DEGs (irinotecan, 97.09). For imatinib, four clinical trials were conducted, with three trials (NCT00080665, NCT00087152, and NCT00193180) indicating no therapeutic effect and one trial (NCT00372476) supporting a treatment effect[61–64]. Consequently, imatinib was deemed to have a therapeutic effect on BRCA. Regarding metformin, two clinical trials were undertaken, and all of them (NCT01101438, NCT01310231) reported opposing treatment effects, with one trial (NCT01310231) even suggesting a worse prognosis with metformin treatment[65, 66]. As for irinotecan, two clinical trials were conducted, both of which supported a treatment effect (NCT00079118, NCT00083148)[67, 68]. In the case of BRCA, PCS accurately predicted the efficacy of one drug and the lack of efficacy of another, while DEGs erroneously predicted a lack of treatment effect for one drug.

In the context of HNSC, two drugs were identified using the PCS method (irinotecan, 69.71; paclitaxel, 83.71). In the case of irinotecan, two clinical trials (NCT00040807, NCT00639769) reported opposing treatment effects, aligning with the correct prediction [69, 70]. For paclitaxel, there were nine clinical trials (NCT00002632, NCT00003193, NCT00005087, NCT00011999, NCT00014118, NCT00095927, NCT00301028, NCT00392704, NCT01084083) supporting a treatment effect, while three clinical trials (NCT00002922, NCT00002888, NCT00006248) did not support a treatment effect, which represents an incorrect prediction by PCS[71–81].

In the context of LIHC, PCS analysis identified eight drugs (everolimus, -99.01; gemcitabine, -92.5; doxorubicin, -86.05; docetaxel, -84.67; ispinesib, -76.64; idarubicin, -63.00; pazopanib, 67.17; pazopanib, 89.83), and two drugs were identified using DEGs analysis (gemcitabine, 88.26; megestrol, 64.11). For everolimus, a single trial (NCT01005199) indicated no treatment effect, resulting in an incorrect prediction by PCS [82]. Conversely, gemcitabine showed consistent treatment effects across all three trials (NCT00006010, NCT00703365, NCT02527772), and PCS analyses correctly predicted this outcome while DEGs not[83–85]. In the case of doxorubicin, where ten clinical trials were conducted, two trials (NCT00003912, NCT00047229) opposed the treatment effect, while eight trials (NCT00012324, NCT00108953, NCT00293397, NCT00471484, NCT00478374, NCT00956930, NCT01445535, NCT02527772) supported it, which suggested a correct prediction by PCS[85–93]. Similarly, docetaxel showed consistent support for treatment effects in both trials (NCT00006010, NCT00532441), affirming the accuracy of PCS prediction [83, 94]. For ispinesib, one trial (NCT00095992) indicated treatment effect, aligning with PCS prediction [95]. Idarubicin also received support for treatment effects from clinical trial NCT02185768[96]. Pazopanib’s treatment effect was affirmed by clinical trial NCT00370513[97]. Lenalidomide was supported by NCT00717756 while megestrol was opposed by NCT00041275 [98, 99]. Consequently, PCS accurately predicted the efficacy of ispinesib and idarubicin, yet it inaccurately predicted the outcomes for pazopanib and lenalidomide. The analysis based on DEGs made an accurate prediction for megestrol.

As both LUAD and LUSC are classified under non-small cell lung cancer, we combined their analysis for discussion. In LUAD, In LUAD, eight drugs were identified using PCS (fostamatinib, -95.75; everolimus, -95.05; doxorubicin, -94.16; dasatinib, -79.01; linifanib, -73.73; dexamethasone, -71.69; enalapril, -65.62; azacitidine, 56.68). The efficacy of fostamatinib was supported by the trial NCT00923481, and a clinical trial, NCT00096486, provided evidence for the treatment effect of everolimus[100]. Three clinical trials (NCT00002822, NCT00003364, NCT00003847) corroborated the treatment effect of doxorubicin[101–103]. For dasatinib, two trials (NCT00787267, NCT00787267) supported treatment effect, and one trial (NCT00570401) opposed it[104, 105]. Linifanib had support from one clinical trial (NCT01225302), while dexamethasone had backing from two trials (NCT00247416, NCT00247416), and enalapril and azacitidine were each supported by one clinical trial (NCT01754909 and NCT00387465, respectively)[106–109]. In LUAD, PCS accurately predicted the efficacy for seven drugs and made one incorrect prediction.

In LUSC, six drugs were identified using PCS (dasatinib, -96.34; fostamatinib, -96.23; iloprost, -81.48; doxorubicin, -64.18; azacitidine, 50.99; disulfiram, 76.62), and one drug was identified using DEGs (entinostat, 77.8). Out of these seven drugs, four drugs (dasatinib, doxorubicin, fostamatinib, and azacitidine) overlapped with those identified in LUAD. For the overlapped four drugs, PCS made accurate predictions for dasatinib, doxorubicin, and fostamatinib, while providing an inaccurate prediction for azacitidine, consistent with the results observed in LUAD. As for the remaining three drugs, each had one trial support for their treatment effect (NCT00084409 for iloprost, NCT00312819 for disulfiram, and NCT00387465 for entinostat)[109–111]. Thus, in LUSC, PCS predicted accurately predicted 4 drugs and made 2 incorrect predictions while DEGs only made one incorrect prediction.

In THCA, two drugs were identified using PCS (sorafenib, 56.31; selumetinib, 97.43). A clinical trial, NCT00984282, supported the treatment effect of sorafenib, while another clinical trial, NCT00559949, opposed the treatment effect of selumetinib[112–114]. Therefore, in THCA, PCS made one correct prediction and one incorrect prediction, respectively.

Subsequently, we compiled the validation outcomes of PCS in Table 4 and DEGs in Table 5, extracted from supplementary Table 2. In PCS, the validation results for drugs were arranged based on their scores, as presented in Table 4. There were 28 validation results in PCS, comprising 18 negative score validations (below -50) and 10 positive score validations (above 50). Examining Table 4, it is evident that among the 18 negative score validation results, 17 were accurately validated, with the exception of everolimus. Among the 10 positive score validation results, 5 were correctly validated. For DEGs, the validation results for drugs were similarly organized by score. Table 5 summarizes 4 validation results, only including 4 positive score validation. Among the 4 positive score validations, 3 were incorrectly predicted and only one was accurately predicted. Given that disease-specific signature-based drug repurposing approaches primarily focus on negative score hits, deemed as candidate drugs, PCS exhibited exceptional performance in drug prediction with high accuracy (17/18), while DEGs got nothing validated. Notably, it was surprising that positive score hits yielded many effective treatment drugs. According to the established principle, negative score drugs exhibit treatment effects, near-zero score drugs have no effect, and positive score drugs promote tumor progression. Consequently, only negative score drugs warrant attention. However, half of the positive score drugs demonstrated treatment effectiveness. It appears that both positive and negative score drugs contribute to enriching treatment-effective drugs, with negative score hits being more prevalent.

### Top 5 predicted drugs for the six types of tumor

Given the robust predictive capability of our PCS signatures-based drug prediction, we investigated the top 5 predicted drugs (i.e., the top 5 compounds with negative scores) for each of the six types of tumors. Additionally, for comparison, we plotted the top 5 positive score compounds. In BRCA, the top 5 negative score compounds comprised amonafide, cobalt(II)-chloride, piperacillin, RO-90-7501, and calyculin (Figure 5A). Notably, amonafide is a drug under investigation for BRCA treatment. Piperacillin, an approved bacterial cell wall synthesis inhibitor, is utilized for treating polymicrobial infections. Cobalt(II)-chloride acts as a heat shock protein (HSP) inducer, while RO-90-7501 is a beta-amyloid inhibitor, and calyculin serves as a protein phosphatase inhibitor. For HNSC, the top 5 negative score compounds included cobalt(II)-chloride, Cefaclor, HNHA, rhapontin, and oleanolic-acid (Figure 5B). As mentioned above, Cobalt(II)-chloride is a heat shock protein(HSP) inducer. Cefaclor, also a bacterial cell wall synthesis inhibitor, an approved drug, is used for the treatment of certain infections caused by bacteria such as pneumonia and ear, lung, skin, throat, and urinary tract infections. HNHA is a histone deacetylase (HDAC) inhibitor. Rhapontin is an apoptosis stimulant. Oleanolic-acid is a G protein-coupled receptor agonist. In LIHC, the top five negative score compounds consisted of BMS-345541, XMD-1150, entinostat, everolimus, and BI-2536 (Figure 5C). Notably, entinostat (an HDAC inhibitor) was under investigation for some disease including breast cancer. Everolimus, an mTOR inhibitor, is an approved drug used for treating advanced hormone receptor-positive, HER2-negative breast cancer. BMS-345541, XMD-1150, and BI-2536 are inhibitors targeting IKK, leucine-rich repeat kinase, and polo-like kinase, respectively. In LUAD, the top 5 compounds were givinostat, TG-101348, ISOX, XMD-892, and XMD-1150 (Figure 5D). Givinostat (an HDAC inhibitor) was under investigation for various conditions, including Polycythemia Vera and Juvenile Idiopathic Arthritis. TG-101348 is a FLT3 inhibitor, ISOX is an HDAC inhibitor, and XMD-892 is a MAP kinase inhibitor. XMD-1150 is a leucine-rich repeat kinase inhibitor. For LUSC, the five compounds were ellipticine, AZD-7762, PD-184352, AS-703026, and mirdametinib (Figure 5E). Ellipticine is a topoisomerase inhibitor with potent antineoplastic properties. AZD-7762 (a CHK inhibitor) is under investigation for various cancers, and mirdametinib (a MEK inhibitor) is studied for melanoma, solid tumors, and breast neoplasms. Both PD-184352 and AS-703026 are MEK inhibitors. In THCA, only one compound with a score lower than -90 and one between -80 to -90 were identified. These two compounds, verrucarin-a and omacetaxine mepesuccinate, are shown in Figure 5F. Omacetaxine mepesuccinate, a protein synthesis inhibitor, is an approved drug for accelerated or chronic phase CML. Verrucarin-a is also a protein synthesis inhibitor.

**Figure 5.**
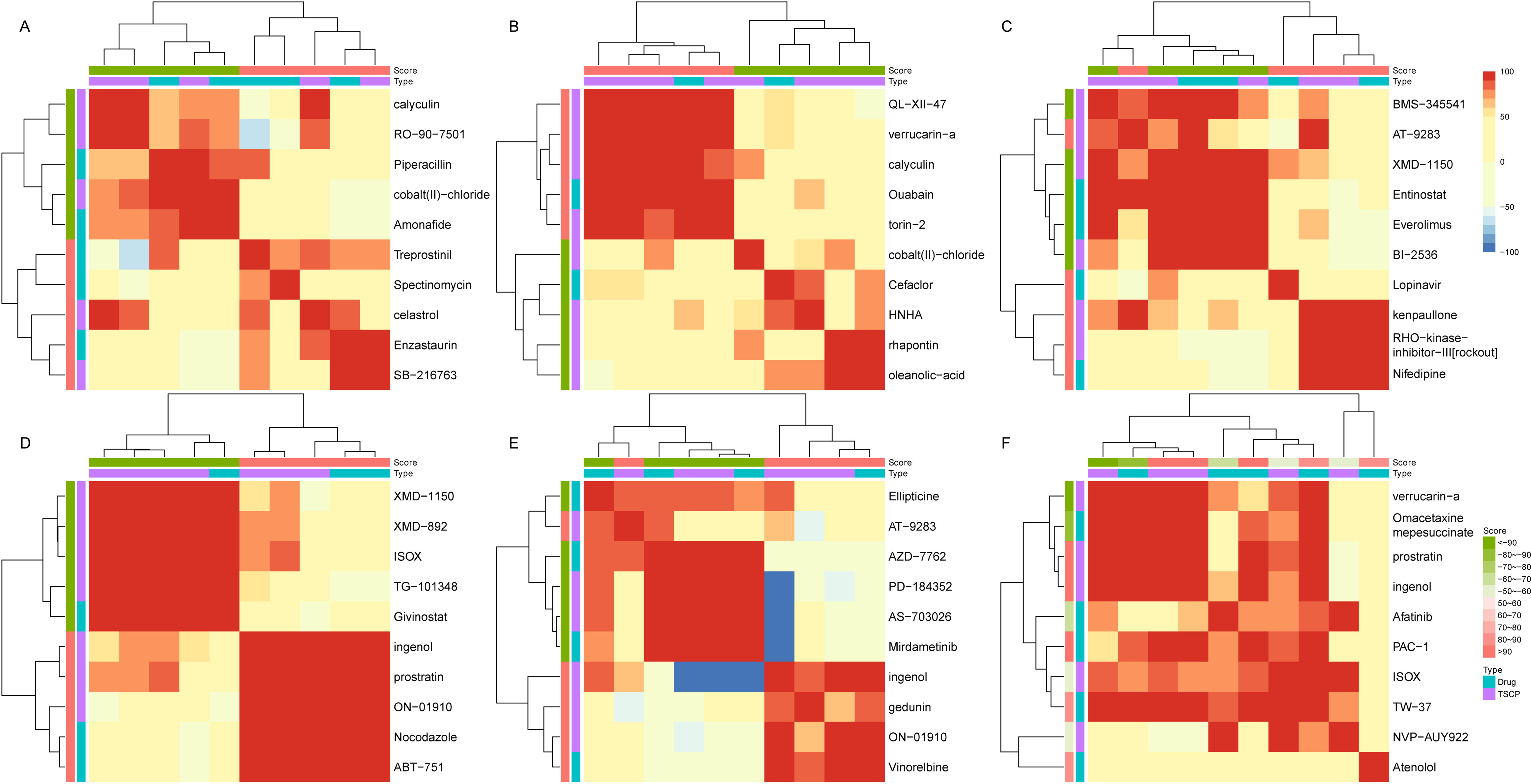
The top five predicted candidate drugs for the six types of tumors. A: BRCA; B: HNSC; C: LIHC; D: LUAD; E: LUSC; F: THCA.

Notably, most of the top five compounds were associated with anti-tumor treatment or shared mechanisms of action with anti-tumor compounds, except for two anti-microbial infection compounds. It is worth mentioning that studies have highlighted the role of intratumoral microbiota in cancer initiation, development, immunity, heterogeneity, therapeutic efficacy, and metastatic colonization.[115–119]. Appropriated antibiotic treatment in tumors has been associated with improved prognosis[115].

### Drugs combination based on PCS signatures

The reliance on a single drug for treating tumors often leads to drug resistance. Addressing this challenge necessitates the identification of appropriate drug combinations. In this study, we sought to discover effective drug combinations for tumor treatment based on PCS signatures, with a focus on BRCA. As illustrated in Figure 4A, we selected four drugs with scores below -50: amonafide, anastrozole, ibuprofen, and vinblastine. The connectivity scores between these drugs are depicted in Figure 6A. Notably, these drugs did not exhibit particularly high similarity to each other. The connectivity score between amonafide and vinblastine was 67.30, while the score between anastrozole and ibuprofen was 50.33. Scores for the other pairs were below 50.

**Figure 6.**
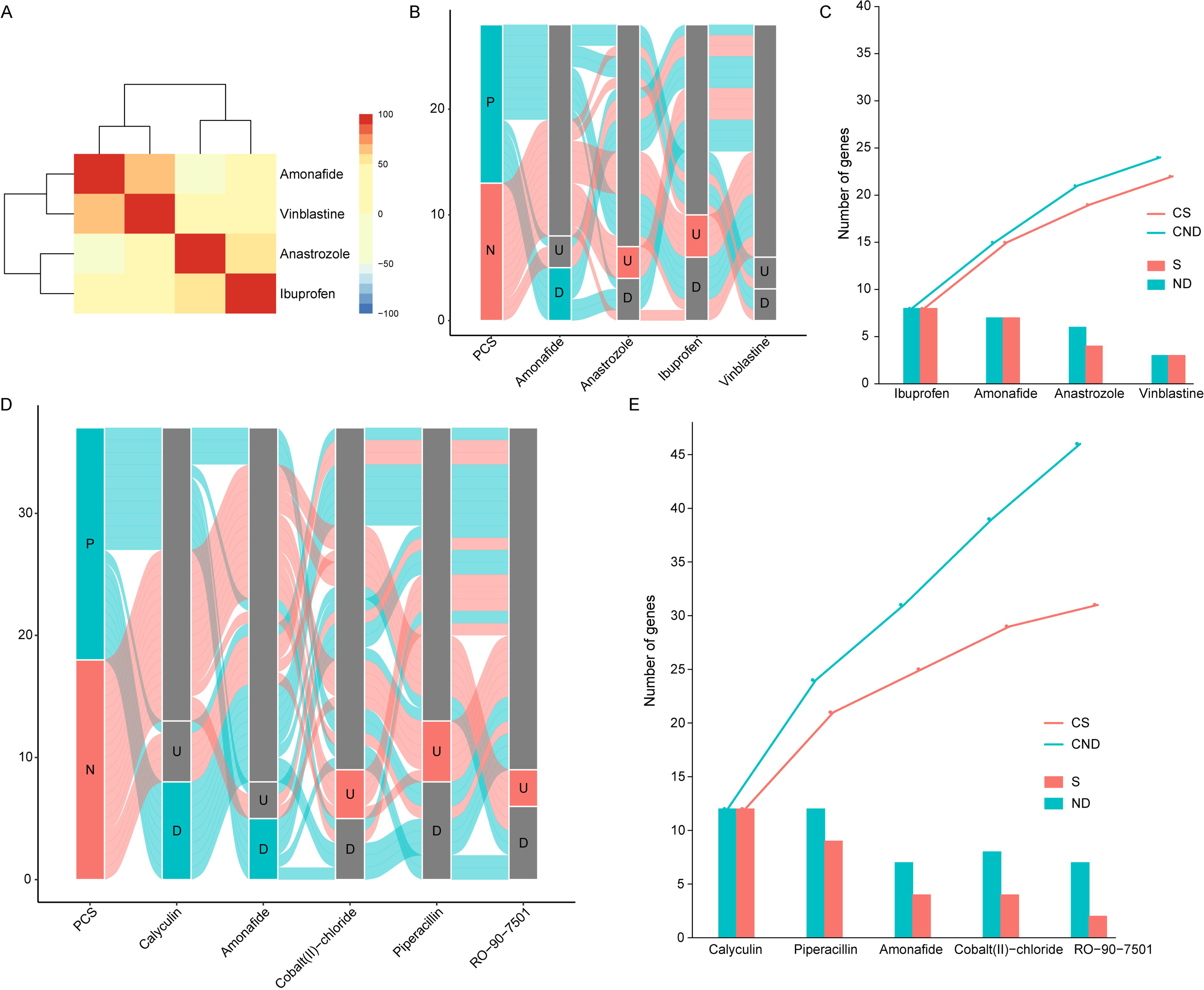
Drug combination order in BRCA for the top four drugs and the top five predicted candidate drugs. A: Connectivity heatmap of the four drugs with score lower than -50, the color bar represented connectivity socore; B: Sankey diagram between PCS based signatures of BRCA and the four drugs, P represented PCS based BRCA signature genes with positive PCS, N represent PCS based BRCA signature genes with negative PCS, U represented up-regulated genes of drugs, D represented down-regulated genes of drugs; C: Drugs combination order of the four drugs, ND represented the overlapped gene number between disease signature and drug signature without set diff (no diff, ND), S represented the overlapped gene number between disease signature and drug signature after set diff (specific, S), CDN represented the cumsum of ND, CS represent the cumsum of S; D: Sankey diagram between PCS based signatures of BRCA and the five predicted candidate drugs; E: Drugs combination order of the five predicted candidate drugs.

Subsequently, we investigated the overlap among the signature genes, as shown in Figures 6B and 6C. Figure 6B indicates that vinblastine demonstrated less favorable characteristics, with a mixture of both up-regulated and down-regulated PCS genes and a minimal gap between the proportions of negative and positive PCS genes. Moreover, Figure 6C highlights that vinblastine had the fewest number of genes. Considering the moderate connectivity score between amonafide and vinblastine, a potential combination strategy could involve amonafide, anastrozole, and ibuprofen. Alternatively, considering the moderate connectivity score between anastrozole and ibuprofen, the combination could consist of amonafide and ibuprofen. Figure 6C illustrates the combination order of drugs based on signature, with ibuprofen being the single-drug scenario, and amonafide and ibuprofen for a two-drug combination.

We applied this methodology to BRCA based on the top negative score compounds shown in Figure 5A. As demonstrated in Figures 6D and 6E, all five compounds exhibited promising characteristics. When combined based on the order of specific genes through step-wise set difference, the optimal combination order was calyculin, piperacillin, amonafide, cobalt(II)-chloride, and RO-90-7501, as shown in Figure 6E. It is important to note that our strategy represents an exploratory analysis and is not intended for treatment recommendations or advice.

### Specific of the PCS based signature genes

As previously outlined, we selected the top and bottom 150 genes ordered by PCS or logFC, respectively, to serve as tumor-specific signatures. Subsequently, we examined their specificity. For each of the six types of tumors, there were 300 genes, resulting in a theoretical total of 1800 genes if every gene were tumor-specific. Upon merging the genes in the PCS signature, a total of 1619 genes were identified, indicating a specificity of 89.94%, as illustrated in Figure 7A and 7B. In contrast, upon merging the six signatures in DEGs, only 1056 genes were identified, accounting for a specificity of 58.67%, as depicted in Figure 7C and 7D.

**Figure 7.**
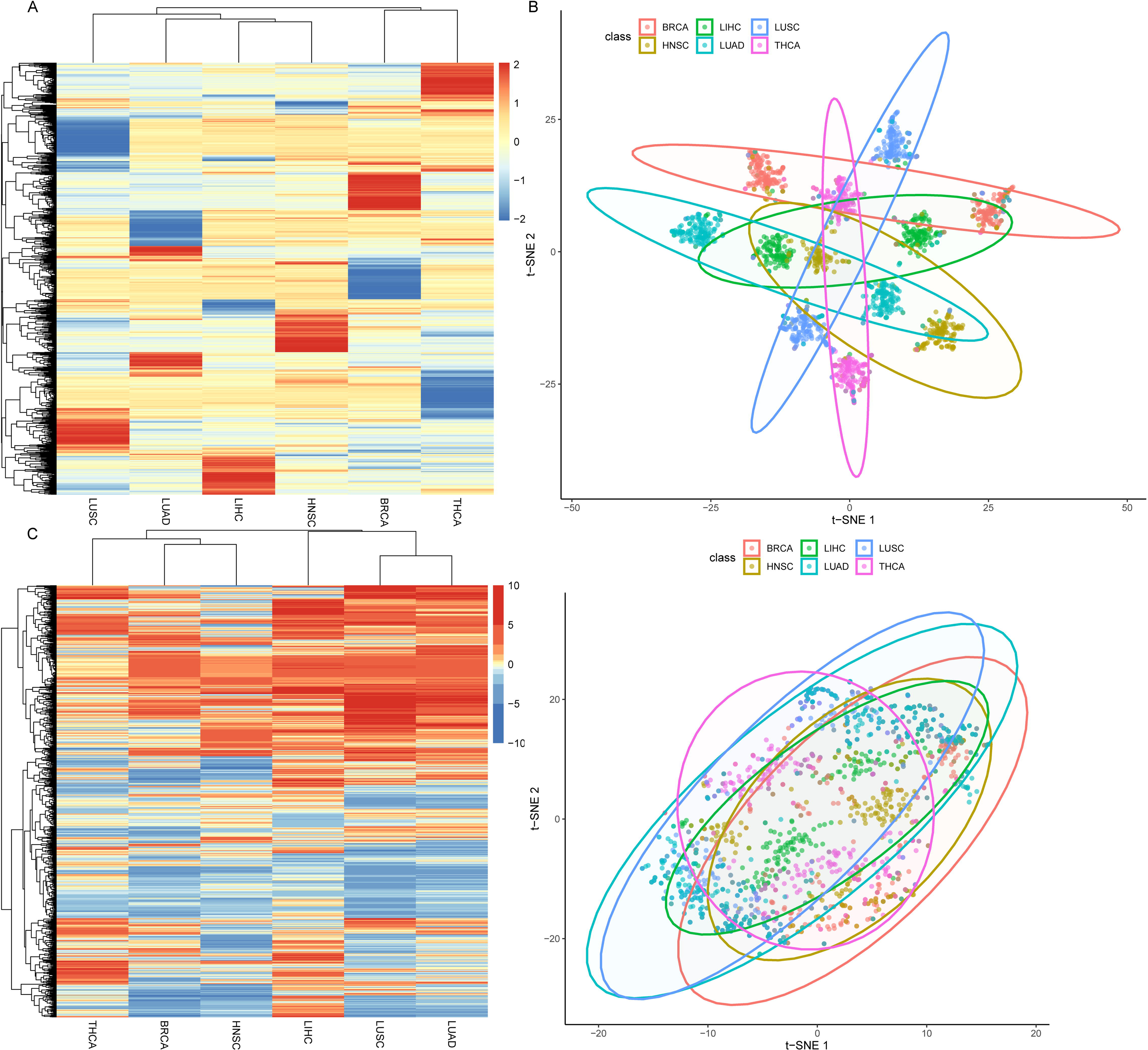
Disease specificity of PCS and DEGs based disease signatures. A: Heatmap of PCS signature genes in the six types of tumor, the color bar represented PCS; B: T-sne map the PCS signatures; C: Heatmap of DEGs signature genes in the six types of tumor, the color bar represented loGFC; D: T-sne map the DEGs signatures.

Figure 7A and 7B highlight that each type of tumor exhibited distinct gene modules in the PCS signature. However, Figures 7C and 7D suggest a lack of discernible specific gene modules in DEGs. Fisher’s exact test revealed that PCS-based signatures were significantly more specific than those derived from DEGs (*ratio* = 6.30, *P*-value < 0.001).

### Association between disease related genes and PCS

Given the high specificity of PCS-based signature genes observed in the preceding results, we investigated the association between DRG and PCS. DRGs and DGS were obtained from the CURATED subset of the DisGeNET database.

In BRCA, for PCS, as depicted in Figure 8A, genes were ranked by PCS, and BRCA-related genes were plotted based on their ranks and DGS. The density distribution of BRCA-related genes and the sum of DGS for these genes were displayed. Subsequently, the difference in PCS between all genes and DRGs was presented. Both sets of genes were ranked by the absolute value of PCS, and the ranks were scaled to the same range. Student’s t-test was employed to examine the PCS differences between all genes and DRGs. Figure 8A illustrates that DRGs tended to be located at both high and low PCS values, with the sum of DGS following a similar pattern. Although the absolute value of PCS for DRGs was slightly higher than that of all genes, the difference was not statistically significant (t-test, *P*-value = 0.15). For DEGs, as shown in Figure 8B, DRGs appeared to be evenly distributed along the rank. The absolute value of logFC in DRGs was higher than that in all genes (t-test, *P*-value < 0.001). To assess the difference in DRG enrichment between PCS and DEGs, all genes were ranked by the absolute value of PCS or logFC. DRGs were then divided into two groups based on the median rank for both PCS and logFC. Fisher’s exact test was employed to evaluate the differences. In BRCA, Fisher’s exact test revealed that PCS exhibited a higher enrichment power for DRGs (*ratio* = 1.20, *P*-value = 0.02). Similar results were observed in HNSC (*ratio* = 1.78, *P*-value = 0.006), LIHC (*ratio* = 1.30, *P*-value = 0.01), LUSC (*ratio* = 1.45, *P*-value < 0.001), and THCA (*ratio* = 1.62, *P*-value = 0.02), as presented in Supplementary Figures 1. Conversely, in LUAD, PCS and DEGs showed comparable results (*ratio* = 1.16, *P*-value = 0.15), as illustrated in Supplementary Figures 1G and 1H. In THCA, the PCS of all genes exhibited a higher value than that of DRGs (t-test, *P*-value = 0.05), whereas for DEGs, there was no significant difference (t-test, *P*-value = 0.83) (Supplementary Figure 1I and 1J).

**Figure 8.**
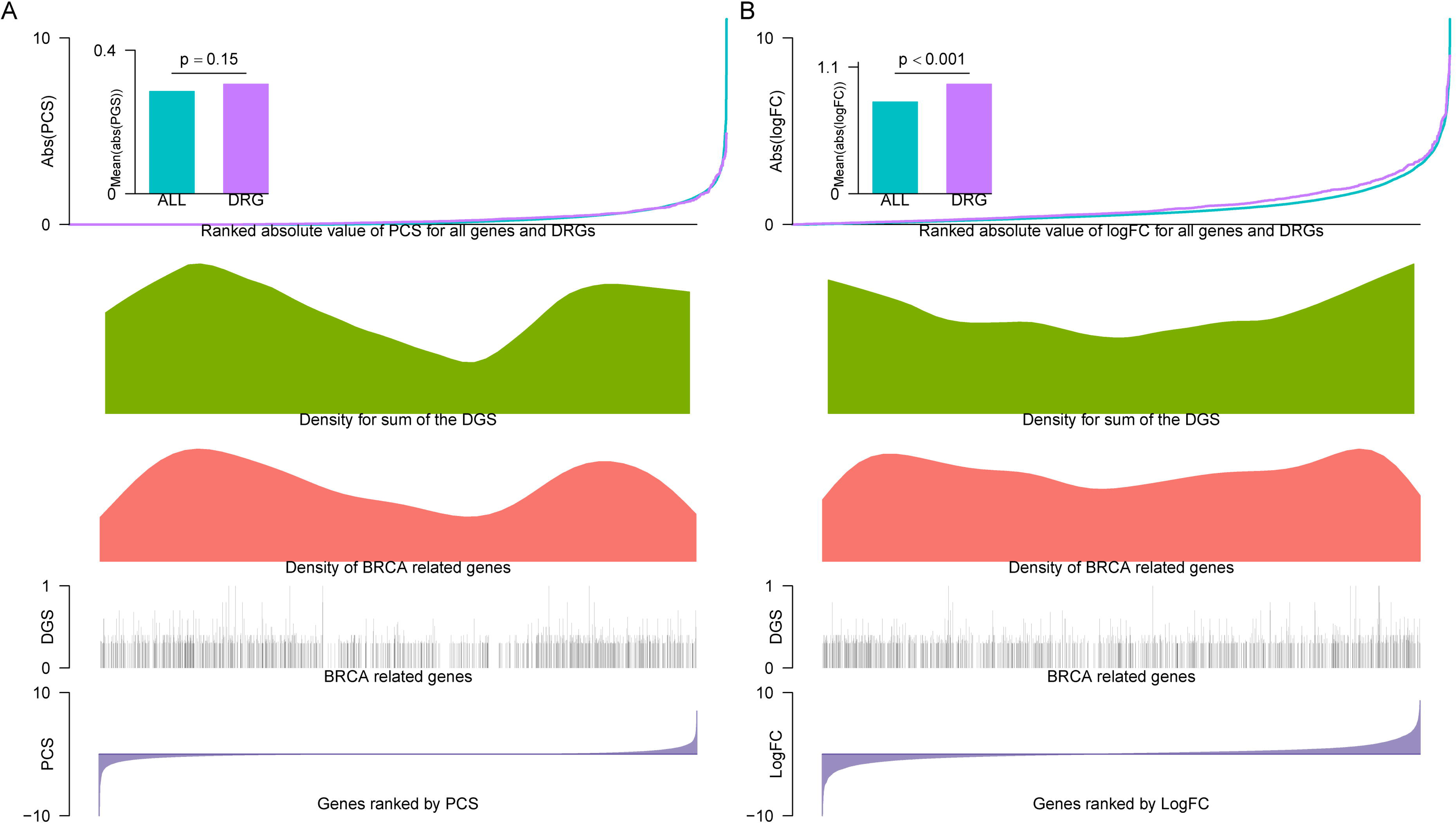
Association between disease related genes and PCS or DEGs in BRCA. DGS: Disease gene score; DRG: Disease related genes.

Surprisingly, Fisher’s exact test indicated that PCS also demonstrated superior enrichment for DRGs compared to DEGs (*ratio* = 1.62, *P*-value = 0.02). This observation prompted a puzzling contradiction observed in both BRCA and THCA. Despite no discernible difference in PCS between all genes and DRGs, with PCS of all genes even surpassing that of DEGs, DEGs displayed better results. However, the enrichment power for DRGs of PCS remained higher.

We hypothesize two plausible explanations for this discrepancy. First, the absolute value of logFC in DEGs was higher than that in PCS. Consequently, DEGs encompassed more genes exhibiting elevated logFC compared to PCS. Second, PCS demonstrated enrichment for DRGs across a spectrum of values, encompassing both extremely high and low values. Irrespective of the nuances between all genes and DRGs, PCS exhibited superior performance in DRG enrichment across five of the six tumor types.

## Discussion

The process of new drug discovery is characterized by its time-intensive, costly, and high-risk nature. Addressing these challenges necessitates effective strategies such as drug repurposing and combination therapies[120–122]. The Cmap approach, based on signature similarity, is a widely used method in drug repurposing. However, the commonly employed method for signature construction involves the use of DEGs. This approach overlooks the consistency between dysregulated genes and their prognostic implications, as well as the concordance of gene roles in prognosis between tumor and normal tissues. Consequently, it may lead to inaccurate or even harmful drug predictions. To overcome these limitations, we introduced a new method termed PCS in this study, which takes into account the aforementioned consistencies. Comparing PCS-based signatures with DEG-based signatures, we observed that PCS was capable of identifying more drugs and exhibited higher prediction accuracy, as validated by DrugBank annotations. To support the predicted drugs lacking corresponding indications, the ClinicalTrials database was consulted, revealing that the majority of predicted drugs were substantiated in clinical trials. Intriguingly, a significant proportion of predicted harmful drugs also demonstrated treatment efficacy. We attempted to propose a drug combination strategy based on the overlap between disease and drug signatures. Ultimately, PCS-based signatures demonstrated high disease specificity and an association with disease-related genes. However, several limitations should be acknowledged. Firstly, the availability of tumor-normal paired samples with RFS was insufficient, with a maximum of 53 pairs across tumor types, excludingBRCA. Secondly, our method lacked comparisons with alternative approaches beyond DEGs. Thirdly, PCS exhibited limited predictive utility in certain tumor types, suggesting its potential unsuitability for all solid tumors. Lastly, PCS had a notable flaw, as positive scores also enriched some treatment drugs, and even predicted harmful drugs were proven to have treatment effects. Although PCS outperformed DEGs, there is still room for refinement and improvement.

**Table 4.** Validation of PCS predicted candidate drugs via ClinicalTrials.gov.

| Drug | Score | Indication | Tumor | Drug | Score | Indication | Tumor |
| --- | --- | --- | --- | --- | --- | --- | --- |
| everolimus | -99.01 | No | LIHC | enalapril | -65.62 | Yes | LUAD |
| dasatinib | -96.34 | Yes | LUSC | doxorubicin | -64.18 | Yes | LUSC |
| fostamatinib | -96.23 | Yes | LUSC | idarubicin | -63.00 | Yes | LIHC |
| fostamatinib | -95.75 | Yes | LUAD | imatinib | -51.34 | Yes | BRCA |
| everolimus | -95.05 | Yes | LUAD | azacitidine | 50.99 | No | LUSC |
| doxorubicin | -94.16 | Yes | LUAD | metformin | 55.28 | No | BRCA |
| gemcitabine | -92.50 | Yes | LIHC | sorafenib | 56.31 | Yes | THCA |
| doxorubicin | -86.05 | Yes | LIHC | azacitidine | 56.68 | Yes | LUAD |
| docetaxel | -84.67 | Yes | LIHC | pazopanib | 67.17 | Yes | LIHC |
| iloprost | -81.48 | Yes | LUSC | irinotecan | 69.71 | No | HNSC |
| dasatinib | -79.01 | Yes | LUAD | disulfiram | 76.62 | No | LUSC |
| ispinesib | -76.64 | Yes | LIHC | paclitaxel | 83.71 | Yes | HNSC |
| linifanib | -73.73 | Yes | LUAD | lenalidomide | 89.83 | Yes | LIHC |
| dexamethasone | -71.69 | Yes | LUAD | selumetinib | 97.43 | No | THCA |

**Table 5.** Validation of DEGs predicted candidate drugs via ClinicalTrials.gov.

| Drug | Score | Indication | Tumor | Drug | Score | Indication | Tumor |
| --- | --- | --- | --- | --- | --- | --- | --- |
| gemcitabine | -88.26 | 1 | LIHC | megestrol | -64.11 | 0 | LIHC |
| entinostat | -77.8 | 1 | LUSC | irinotecan | 97.09 | 1 | BRCA |

## Conclusions

This study introduced PCS as a novel disease-specific signature building method for drug repurposing via the Connectivity Map. PCS effectively addressed the inconsistencies between dysregulated genes and their prognostic implications in tumor tissue, as well as the disparities in prognostic gene roles between tumor and normal tissues. The results demonstrated that PCS outperformed DEGs in drug repurposing predictions and exhibited higher disease specificity. Furthermore, PCS displayed associations with disease-related genes, emphasizing its potential as a valuable tool in the field of drug repurposing.

## List of abbreviations

BRCA: Breast invasive carcinoma
CLUE: Cmap and Lincs Unified Environment
Cmap: Connectivity Map
DEGs: Differential expressed genes
DGS: Disease gene score
DRG: Disease-related genes
FDR: False discovery rates
GWAS: Genome-wide association studies
HNSC: Head and neck squamous cell carcinoma
HSP: Heat shock protein
LIHC: Lung adenocarcinoma
LINCS-L1000: Library of Integrated Network-Based Cellular Signatures L1000
LUAD: Lung adenocarcinoma
LUSC: Lung squamous cell carcinoma
MODZ: Moderated z-score
NN: Normal Cox negative coefficient
NP: Normal Cox positive coefficient
PCS: Prognosis consistency score
PPI: Protein-protein interaction
RFS: Relapse-free survival
RNA-seq: RNA-sequencing
TCGA: The Cancer Genome Atlas
TD: Tumor down-regulated genes set
THCA: Thyroid carcinoma
TN: Tumor Cox negative coefficient
TP: Tumor RFS Cox positive coefficient
TPM: Transcripts per million
TRFSN: Tumor RFS Cox negative coefficient set
TRFSP: Tumor RFS Cox positive coefficient set
TU: Tumor up-regulated genes set
TWAS: Transcriptome-wide association studies
UCRDP: Unconsistency ratio of dysregulation and RFS prognosis
UCRTPNP: Unconsistency ratio between tumor RFS prognosis and normal RFS prognosis.

## Declarations

### Data Availability statement

All data used in this study were open access.

### Authors’ contributions

Wei Ma conceived and designed the study. Jun Li and Ming Xiong supervised the implemented the study. Mingyue Hao collected the data and performed some computational analysis. Dandan Li and Yuanyuan Qiao performed some validation analysis and drafted the manuscript. Wei Ma and Mingyue Hao plotted the figures. Wei Ma, Jun Li and Ming Xiong revised the manuscirpt.

---

**Supplementary Figure 1. Association between disease related genes and PCS or DEGs in HNSC, LIHC, LUSC, LUAD, and THCA.**

DGS: Disease gene score; DRG: Disease related genes.

**Supplementary table 1. Summary of 190 articles.**

**Supplementary table 2. Validation detail of predicted candidate drugs via ClinicalTrials database.**

