## Supplementary figures and images for "Tumor relapse-free survival prognosis related consistency between cancer tissue and adjacent normal tissue in drug repurposing for solid tumor via connectivity map"

### Supplementary figure 1

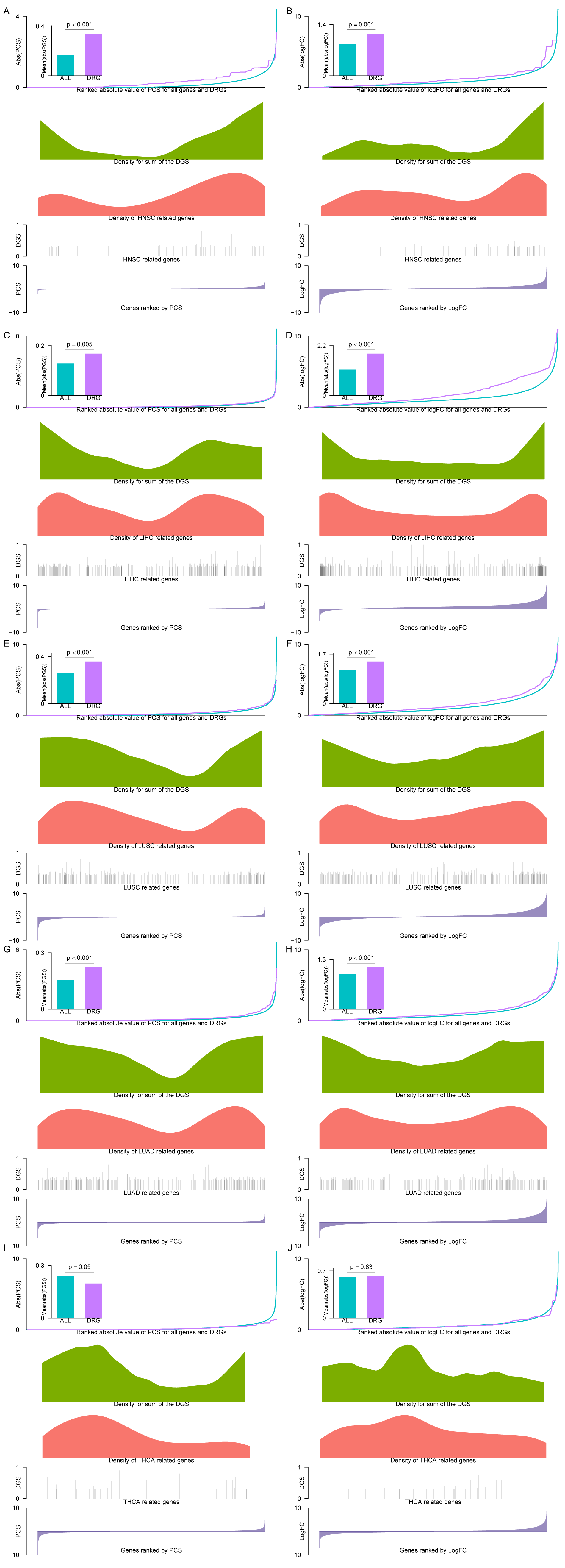
